# Asymmetric spin echo multi-echo echo planar imaging (ASEME-EPI) sequence for pre-clinical high-field fMRI

**DOI:** 10.1101/2024.10.12.617985

**Authors:** Kyle A. Johnson, Christopher P. Pawela, Andrew S. Nencka, Jason W. Sidabras

## Abstract

In functional magnetic resonance imaging (fMRI) of the blood oxygen level-dependent (BOLD) contrast, gradient-recalled echo (GRE) acquisitions offer high sensitivity but suffer from susceptibility-induced signal loss and lack specificity to microvasculature. In contrast, spin echo (SE) acquisitions provide improved specificity at the cost of reduced sensitivity. This study introduces Asymmetric Spin Echo Multi-Echo Echo Planar Imaging (ASEME-EPI), a technique designed to combine the benefits of both GRE and SE for high-field preclinical fMRI. ASEME-EPI employs a spin echo readout followed by two asymmetric spin echo (ASE) GRE readouts, providing an initial T2-weighted SE image and subsequent T2^*\**^-weighted ASE images. A feasibility study for the technique was implemented on a 9.4 T pre-clinical MRI system and tested using a visual stimulation in northern tree shrews. Comparing ASEME-EPI with conventional GRE echo planar imaging (GRE-EPI) and SE echo planar imaging (SE-EPI) acquisitions, results showed that ASEME-EPI achieved BOLD contrast-to-noise ratio (CNR) comparable to GRE-EPI while offering improved specificity in activation maps. ASEME-EPI activation was more confined to the primary visual cortex (V1), unlike GRE-EPI which showed activation extending beyond anatomical boundaries. Additionally, ASEME-EPI demonstrated the ability to recover signal in areas of severe field inhomogeneity where GRE-EPI suffered from signal loss. The performance of ASEME-EPI is attributed to its multi-echo nature, allowing for SNR-optimized combination of echoes, effectively denoising the data. The inclusion of the initial SE also contributes to signal recovery in areas prone to susceptibility artifacts. This feasibility study demonstrates the potential of ASEME-EPI for high-field pre-clinical fMRI, offering a promising compromise between GRE sensitivity and SE specificity while addressing challenges of T2^*\**^ decay at high field strengths.

## Introduction

Functional magnetic resonance imaging (fMRI) is a powerful technique to study brain function. fMRI detects subtle changes in signal intensity evoked by brain activity [1]. After a stimulus, blood flow to the active region of the brain increases at a rate that out-paces the metabolic rate of oxygen consumption, yielding an increase in blood volume and a dilution of deoxygenated blood [1]. Oxygenated hemoglobin, unlike deoxygenated hemoglobin, is diamagnetic, and oxygenated blood thus has a smaller magnetic susceptibility, *χ*, than deoxygenated blood [2]. The reduced magnetic susceptibility difference between oxygenated blood and surrounding tissue during activation results in an increased T2^*\**^ which yields increased signal intensity in T2^*\**^-weighted images compared to equilibrium [1], [2]. This blood oxygen level dependent (BOLD) contrast [2] is an indirect measure of neuronal activity and is often cited as a quantitative measure to assess brain function [1], [3].

Animal fMRI models allow researchers to evaluate brain function in both typical and pathological conditions with reduced subject-specific confounds which impact human fMRI studies. Further, the smaller required bore diameter for pre-clinical imaging makes high-field imaging more accessible. Pre-clinical fMRI studies typically involve gradient-recalled echo (GRE) experiments to capture T2^*\**^-weighted images for BOLD contrast [4], though ultra-high magnetic field strengths have been used with spin echo (SE) acquisitions to achieve enhanced spatial localization with T2-weighted BOLD contrast [5].

In fact, GRE and SE have distinct sensitivity and specificity profiles which directly affect susceptibility-based contrasts like BOLD [6]. For a fixed voxel volume, Boxerman and colleagues showed GRE, for an intra-vascular magnetic susceptibility difference Δ*χ* on par with a BOLD-driven change in blood oxygenation, has a greater transverse relaxation rate compared to SE for all vascular radii [6]. In addition, GRE has a transverse relaxation rate (ΔR2^*\**^) which peaks for larger vessels like pial veins, compared to a SE acquisition which has a transverse relaxation rate (ΔR2) which peaks for smaller diameter vessels like those found in venules and the capillary bed [6]. This results in extremely sensitive BOLD activation mapping with GRE because larger vessels like pial veins account for greater voxel partial volume fractions than smaller vessels. For GRE, ΔR2^*\**^ peaking for macrovasculature yields activation maps heavily weighted towards large veins which can be spatially distant from neuronal activity [7]. Unlike GRE, SE is uniquely sensitive to the microvasculature, though for an echo time (TE) whereby SE is maximally sensitive, and thus specific, to capillaries, BOLD contrast is approximately 3*×* lower than GRE for a fixed voxel volume and static magnetic field strength *B*_0_ [6].

BOLD vascular sensitivity from Δ*χ* can be altered by deviating from GRE and SE acquisitions through the use of asymmetric spin echo (ASE) acquisitions [6]. ASE imaging involves offsetting the collection of the center of k-space by some time *τ* such that it is temporally shifted relative to the center of the spin echo [6], [8]. As *τ* is increased, the transverse relaxation rate as a function of vessel size curve shifts [6] toward larger vessel sizes which corresponds to changing the vessel size at which the relaxivity curve peaks. At the limit that *τ* becomes large enough, the relaxivity curve approaches that of a GRE acquisition [6], where sensitivity increases but losing specificity of vessel size.

Thus, fMRI acquisitions face a compromise between spatial specificity and contrast-based sensitivity which varies between GRE and SE acquisitions. In summary, SE sequences offer better specificity in detecting the precise location of neural activity, while GRE sequences provide higher sensitivity to small changes in blood oxygenation levels. In both modalities, and with ASE, choice of echo time further impacts the trade-off between detecting subtle brain activation patterns and maintaining strong overall sensitivity to the BOLD response.

Moving beyond these compromises, Posse *et al*. [9] *established that the* contrast-to-noise ratio (CNR) of fMRI experiments benefit from sampling multiple TEs. They found BOLD CNR is maximized when echo times were sampled up to 3.2 *×* T2^*\**^. Their work indicated that a T2^*\**^-weighted averaging of multiple GRE images acquired with different echo times theoretically produces a 30% BOLD CNR increase compared to simple summation, and is dependent on the number echoes [9]. BOLD CNR is maximized when three echoes are included in the weighted summation, plateauing for more than 3 echoes [9].

Given such benefits, multi-echo acquisitions have grown in popularity in human fMRI research [10]. It is the goal of this work to bring such benefits of multi-echo acquisitions to pre-clinical fMRI at high field. However, simple extension of multi-echo echo planar imaging (EPI) to high-field imaging is not practical. An increase in magnetic field strength causes a shortening of T2^*′*^ [11], which results in less available time to acquire multiple images following an excitation. Furthermore, at 9.4 T, high-density channel arrays in pre-clinical fMRI is limited due to the small size of a small mammal brain, so parallel imaging techniques can only marginally reduce readout duration and add additional EPI readouts. Other image acceleration techniques such as Partial Fourier and the use of higher receiver bandwidths provide minimal benefit [12].

To address a shortening T2^*′*^ and gain microvascular sensitivity, a multi-echo acquisition with spin echoes can be used. Some work has been performed to achieve the benefits of spin echo and gradient recalled echo acquisitions with the aptly named SAGE (spin and gradient recalled echo) pulse sequence [13]. While originally developed for quantifying contrast-enhanced perfusion, it has been recently extended to support fMRI studies at 3 T [14]. However, the shortened T2^*′*^ and modest acceleration available in high-field pre-clinical imaging present a barrier to directly port SAGE to high-field fMRI work.

Given the benefits of complimentary sensitivities and specificities of GRE and SE acquisitions, the benefit of multi-echo acquisitions for enhancing BOLD sensitivity, and the limitations of multi-echo GRE and SAGE acquisitions when T2^*′*^ is reduced at higher fields, the purpose of our work was to develop a new multi-echo acquisition for use in small mammal fMRI at high field strengths. Here we introduce the asymmetric spin echo multi-echo echo planar imaging (ASEME-EPI) acquisition at 9.4 T and provide a feasibility study to demonstrate its use in mapping the cortical response to a visual stimulus in a small cohort of northern tree shrews. In this work, we show that ASEME-EPI abides by fundamental susceptibility-contrast principles, has BOLD CNR which approaches that of GRE-EPI, recovers signal in areas of severe field inhomogeneity through the inclusion of the SE, and produces activation maps with cortical activations which do not cross anatomical boundaries that GRE-based activations cross.

## Materials and methods

### ASEME-EPI acquisition

As noted, rapid T2^*′*^ decay presents a challenge for multi-echo acquisitions at high-field. For a high-resolution EPI acquisition with partial Fourier acceleration, the readout can take over 15 ms. If GRE acquisitions are used, the acquired signal can decay to effectively noise by the time a third echo image would be acquired. Similarly, with an acquisition between the excitation and refocusing pulses of the SAGE pulse sequence, T2 decay associated with the duration of the readout and the signal loss from T2 decay and T2^*′*^ decay make the acquisition of more than two images impractical. With these noted limitations, the ASEME-EPI sequence was designed to maximize signal intensity for images with three different contrasts following a radio frequency (RF) preparation.

The ASEME-EPI pulse sequence, shown in Fig 1, begins with a 90*°* and 180*°* RF pulse preparation separated by a time 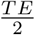. Unlike SAGE, ASEME does not have an image readout between the RF pulses. This allows the RF pulses to be placed close together, allowing a spin echo acquisition to be completed at its minimal achievable time. This spin echo acquisition, with intensity noted as *S*_0,*eff*_, recovers signal loss from regions of extreme field inhomogeneity, and provides BOLD contrast from its T2 weighting alone. Subsequent EPI readouts are two asymmetric spin echo acquisitions with enhanced T2^*′*^ weighting. With this parameterization, the signal decay curve from *S*_0,*eff*_ to the first asymmetric spin echo *S*(*ASE*_1_) and second asymmetric spin echo *S*(*ASE*_2_) is defined by T2^*\**^. Because the spin echo acquisition is acquired with an effective GRE TE of 0 ms, this organization allows the asymmetric spin echoes to have effective GRE echo times which are less than achievable in a pure GRE-based multi-echo acquisition. Additionally, with no acquisition between the excitation and refocusing pulses, the second ASE acquisition (third echo image) is acquired with less T2 and T2^*′*^ decay than the third echo image of a SAGE acquisition, or GRE-based multi-echo acquisition.

**Fig 1.**
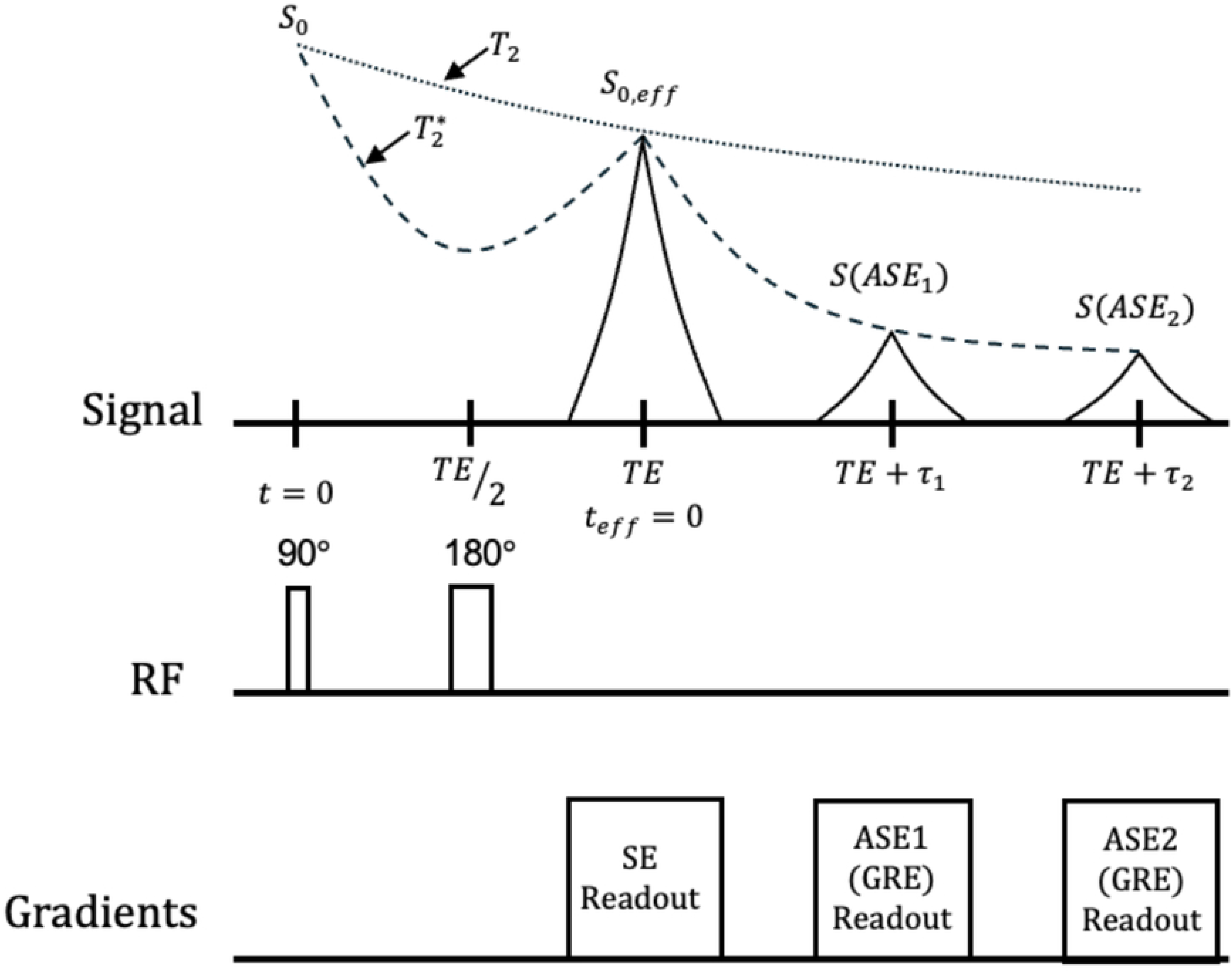
Signal evolution illustration for the ASEME-EPI acquisition. Three echoes are drawn preceded by a 90*°* and 180*°* RF preparation. The first echo is a SE with the seceding echoes being ASEs; the ASEs are GRE readouts. T2^*\**^ and T2 relaxation curves are included. The center of the spin echo occurs at *t*_*eff*_ = 0 which serves as the shifted time origin for analysis. The initial SE signal at *t*_*eff*_ = 0 is *S*_0,*eff*_ with the resultant signal intensities measured for the ASEs having undergone T2^*\**^ attenuation relative to *S*_0,*eff*_. The initial SE signal *S*_0,*eff*_ can be related back to *S*_0_ through T2 decay.

The measured signal intensities *S*_0,*eff*_, *S*(*ASE*_1_), and *S*(*ASE*_2_) are repeated samples of the T2^*\**^ curve, enabling T2^*\**^ estimation. While canonical multi-echo acquisitions typically use *≥* 3 GRE readouts for T2^*\**^ estimation [9], ultra-high field strengths significantly shorten T2^*\**^, resulting in inadequate signal-to-noise ratio (SNR) for longer echo times [2]. ASEME-EPI circumvents this issue by beginning with a SE acquisition which provides the first echo at *t*_*eff*_ of zero. The inclusion of a SE prior to the ASEs significantly improves the combined SNR by providing a signal before T2^*\**^ decay. Therefore, an ASE from an ASEME-EPI acquisition will always have greater SNR than a GRE from a typical multi-echo acquisition at identical echo times. Moreover, an ASE can have greater SNR than a GRE even at different echo times, provided the inequality in Eq (1) is satisfied (derivation is in S1 Appendix).

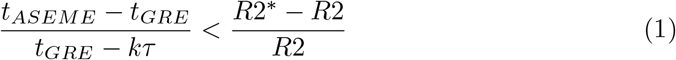

Here, *t*_*ASEME*_, *t*_*GRE*_, and k are the ASEME-EPI echo time, GRE echo time, and echo number, respectively. The integer k represents the chronological echo number in the ASEME-EPI acquisition and can take the values 0 … *n −* 1. The ASEs will have echo numbers *>* 0.

For a three-echo ASEME-EPI acquisition, as shown in Fig 1, ASE2 can have greater SNR than the third echo of an analogous multi-echo acquisition, leading to more accurate T2^*\**^ estimates. By shortening the effective echo time and eliminating the readout between the 90*°* and 180*°* pulses, ASEME-EPI yields higher SNR T2^*\**^ samples and allows for more T2^*\**^ samples. Therefore, an ASEME-EPI sequence will improve T2^*\**^ estimation and provide denoising capabilities for fMRI despite the shortening of T2^*\**^ at high-fields.

### Animal imaging

All work including animals were performed in accordance with an Animal Use Application (AUA2470) approved by the Medical College of Wisconsin’s Institutional Animal Care and Use Committee (IACUC). Three northern tree shrews (*Tupaia belangeri*) were acquired from the Max Planck Florida Institute for Neuroscience. Prior to each imaging session, tree shrews were placed in an induction chamber with 5% isoflurane. Once anesthetized, the tree shrew was inserted into a nosecone whereby isoflurane could be continuously delivered. An intravenous catheter was placed in the saphenous vein to administer dexmedetomidine (0.2 mg/kg/hr) while in the magnet. Dexmedetomidine was infused at a rate of 0.66 mL/hr for the duration of the imaging session in combination with 1% isoflurane. Respiratory rate and temperature were measured via a pneumatic pillow sensor and rectal thermistor probe interfaced to a computer for continuous monitoring.

Experiments were performed on a Bruker BioSpec 94/20 USR 9.4 T pre-clinical magnet. A Bruker 86 mm quadrature volume coil and an in-house built self-resonant spiral surface coil [15] were the transmit and receive channels, respectively. The self-resonant spiral surface coil geometry was designed to maximize SNR within the brain of small mammals and was coupled with an on-board low-noise amplifier (LNA) for further signal improvement. The fMRI studies were conducted using three subjects with the following acquisitions: 1) GRE-EPI, 2) SE-EPI, and 3) ASEME-EPI. Two SE-EPI acquisitions were collected, one to match the TE of the ASEME-EPI SE readout (SEASE; 16 ms), and the other to maximize BOLD sensitivity with TE matching blood T2 (SE40; 40 ms). Imaging parameters common amongst the sequences are summarized in Table 1 along with TEs for each acquisition in Table 2.

**Table 1.** Imaging parameters.

|  |  |
| --- | --- |
| Matrix Size | 75 x 75 |
| Field of View (mm) | 30 x 30 |
| Slice Thickness (mm) | 0.4 |
| Number of Slices | 10 |
| Acquisition Bandwidth (kHz) | 250 |
| Number of Overscan Lines | 16 |
| Repetition Time (ms) | 2000 |
| Number of Repetitions | 110 |

**Table 2.** Echo times.

|  |  |
| --- | --- |
| GRE | 18 ms |
| SE | 16, 40 ms |
| ASEME ($\tau = 15$ ms) | 16 (SE), 31 (ASE), 46 (ASE) ms |

Neurovascular responses were evoked via a custom-built visual stimulation device. Two fiber optic arrays placed proximal to the tree shrew’s eyes delivered bilateral visual stimulus. A 5 Hz flashing white light was presented in three 20 s blocks separated by 40 s rest periods.

The ASEME-EPI pulse sequence was implemented on the Bruker 94/20 imaging system (Paravision 6.0.1), and a customized reconstruction algorithm was written in Python to reconstruct low-signal acquisitions with reduced Nyquist ghosting compared to the default Paravision reconstruction (available upon request).

After reconstruction, the individual ASEME-EPI echoes for each echo time series time point were combined into a single image, as described in Eq (2),

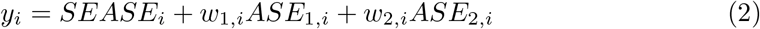

where *y*_*i*_ is the combined image for time point *i, w*_1,*i*_ is the weight used to scale the contribution of the image from the first asymmetric spin echo image for timepoint *i, ASE*_1,*i*_, and *w*_2,*i*_ is the weight for scaling the contribution from the second asymmetric spin echo for timepoint *i, ASE*_2,*i*_.

The weights for the combination were defined following an effective echo time model. First, a voxelwise estimation of T2^*\**^ was performed based upon a task-free acquisition which matched the fMRI acquisition. Low-order (zeroth through second) Legendre polynomials were subtracted from the time series of each echo to account for system drift. The de-trended mean of each echo was calculated and log transformed and a linear regression was performed to estimate R2^*\**^. The estimated R2^*\**^ was then used following Eq (3) to define the weights,

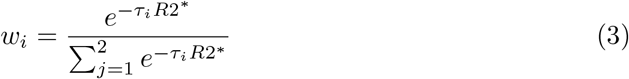

where *τ*_*i*_ is the temporal offset of echo *i* from the spin echo. Thus, the individual time point task images were generated through this weighted averaging, following the idea presented by Posse *et al*. [9].

The experiments in this work used a time relative to *t* = 0 for the GRE acquisition with TE of 18 ms, SE acquisitions with TE of 16 and 40 ms, the ASEME combined acquisition, and each constituent echo of the ASEME acquisition (16, 31, and 46 ms). These time series with matching tasks were analyzed with a General Linear Model (GLM) [16]. For all acquisitions, the GLM consisted of a baseline (*β*_*baseline*_), linear (*β*_*linear*_), and contrast (*β*_*contrast*_) term. The binary stimulus function embedded in the design matrix was temporally shifted by an empirically determined 1 repetition (2000 ms) to account for hemodynamic delay. Voxelwise fits, per acquisition, estimated *β*_*baseline*_, *β*_*linear*_, and *β*_*contrast*_. To account for variance introduced by animal-dependent coil loading and system gains, all time series were scaled such that *β*_*baseline*_ was set to an arbitrary value of 25,000. *β*_*contrast*_ terms and residual time series variance were converted to t-statistics (*H*_0_ : *β*_*contrast*_ = 0, *H*_*a*_ : *β*_*contrast*_ *>* 0).

Fit time series were found to not exhibit independent and identically distributed noise profiles. Thus, an iterative grid search method was employed to determine an appropriate t-statistic threshold, series by series, with matched false positive rates. Voxels outside the brain were subjected to GLM fitting as previously described, and false positive active voxels were defined as those with t-statistics exceeding a specified threshold. Activation t-statistic thresholds were set iteratively using a significance level range of 0.01 to 0.2, with 0.001 increments, resulting in 191 unique t-statistic thresholds. For each threshold, the number of “active” voxels outside of the brain were counted. Statistical thresholds were selected based upon the percentage of these false positive activations outside of the brain. These targets were determined by setting family-wise error (FWE) rates of 0.0005, 0.001, 0.005, 0.01, and 0.05. The empirically determined significance levels for each series and FWE rates are presented in S1 Table.

The empirically determined significance levels for a FWE rate of 0.01 were used to classify active voxels. BOLD CNR was calculated based on the scaled *β* estimates of these active voxels within a mask of the tree shrew brain. Mean CNR, *β*_*contrast*_, and standard deviation of the scaled residuals to the fit model of active voxels per acquisition were reported.

## Results and discussion

Fig 2 depicts the mean contrast-to-noise ratio, contrast, and standard deviation across all active voxels for each series of each animal. Wilcoxon tests do not indicate statistically significant differences between the CNRs of these acquisitions, which vary over a range of approximately 10%. Fig 3 shows maps of active voxels for each acquisition in one animal, with the activations overlaid on the functional imaging time series mean image. Other thresholds are shown in S1 Figure. The plot of CNR shows that acquisitions with more T2^*′*^ weighting (i.e. not spin echo acquisitions) exhibit a trend to higher BOLD contrast. Interestingly, the contrast and standard deviation distributions follow a similar pattern in Fig 2(b) and 2(c), respectively. This is likely a result of the considered active voxels being sensitive to task-related BOLD contrast, which drives the contrast measurements Fig 2(b), while physiologic noise that impacts BOLD contrast beyond task (i.e. respiration volume per time [17], depth of anesthesia, or other vasoactive processes) likely drives the matching pattern in the standard deviation, Fig 2(c).

**Fig 2.**
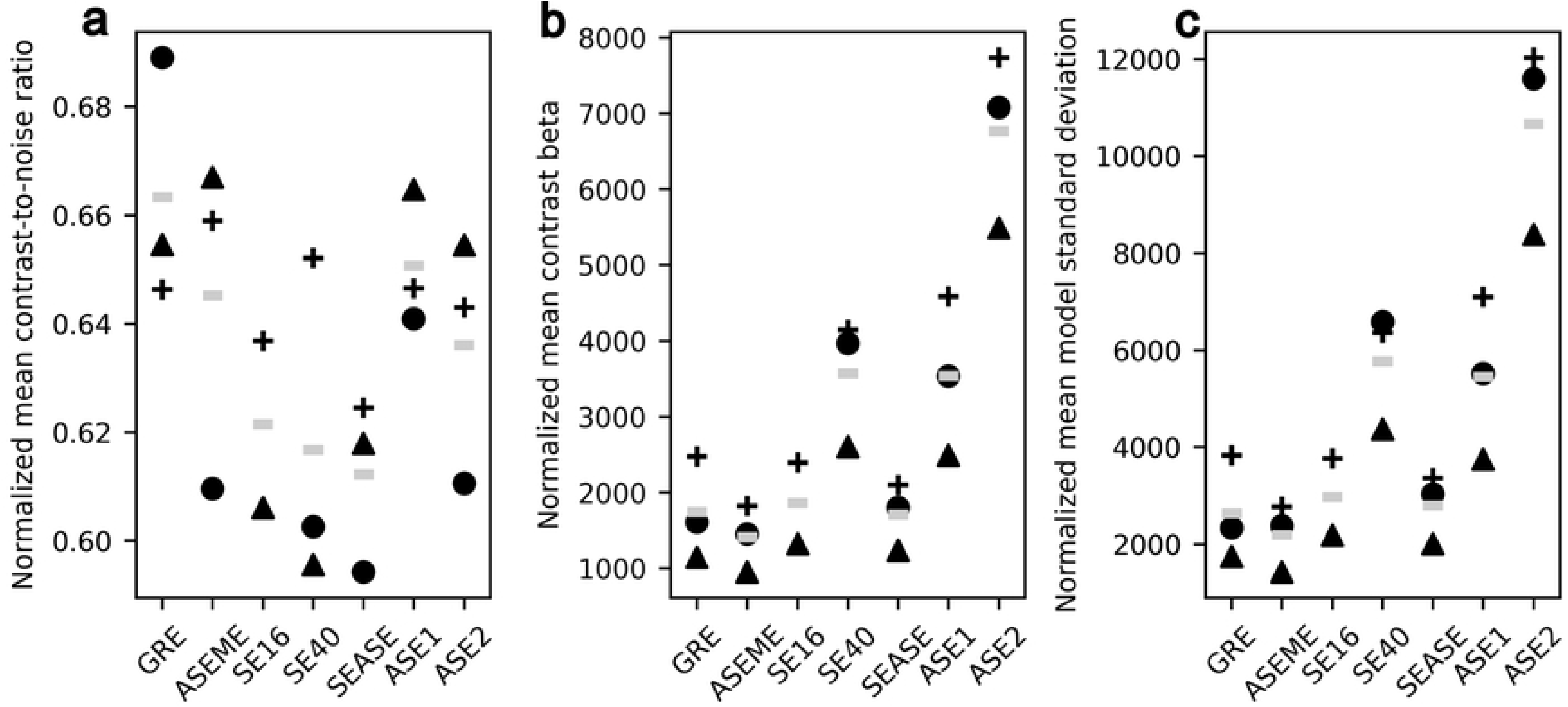
Normalized BOLD mean (a) CNR, (b) *β*_*contrast*_, and (c) standard deviation of the residuals to the fit model. Voxels with t-statistics corresponding to *α ≥ α*_*empirical*_ were plotted; *α*_*empirical*_ values can be found in S1 Table. The shapes designate different subjects and the gray bars mark the inter-subject means. Normalized mean BOLD CNR is normalized *β*_*contrast*_ divided by the normalized standard deviation of the residuals to the fit model, or (b) divided by (c). No significant differences were found between GRE and the other acquisitions after performing a Wilcoxon signed-rank test (*H*_0_ : *µ*_*D*_ = 0, *H*_*a*_ : *µ*_*D*_ *<* 0) on the normalized mean CNRs.

**Fig 3.**
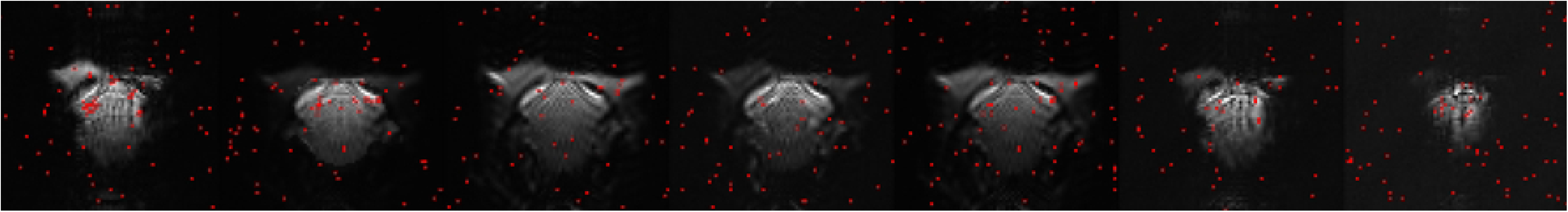
Activation maps in the same slice and the same subject (▴; Fig 2) across the seven acquisitions. T-statistic thresholds were adjusted per acquisition to achieve a FWE of 0.01, and details of the thresholds for each series are provided in S2 Table. Activation maps are shown over EPI images of the respective schemes.

GRE trends to have the highest mean BOLD CNR across all animals, as shown in Fig 2(a), as well as nonspecific activation below V1, seen in column two of Fig 3. GRE exhibiting the largest BOLD CNR is not surprising, it is in concordance with biophysical principles discussed at length in the literature. GRE is ubiquitously sensitive to vessels of all sizes and is greater than SE for all vessels with a radius greater than 6 *µ*m [6].

While GRE offers superior sensitivity, its specificity has been a subject of concern in BOLD literature [18], [19]. The high sensitivity of GRE to vessels of all sizes makes it the typical choice for BOLD imaging, despite its reduced specificity. However, this ubiquitous sensitivity can lead to difficulties in localizing activations to their precise neuronal sources.

Fig 3 illustrates this issue, showing GRE activation that extends beyond the cortical boundary of V1 into adjacent regions [20]. In contrast, ASEME-EPI activation is confined to V1. This difference arises from the underlying spin physics of BOLD responses: GRE’s preference for larger vessels yields activations in displaced locations with larger veins while those large veins create susceptibility gradients that can blur activations across voxels if voxel sizes are smaller than those gradients.

SE acquisitions (SE16, SE40, and SEASE), as shown in Fig 3 and quantified in S2 Table, demonstrate the fewest active voxels among all acquisitions in this study, exhibiting sparse activation maps with small volumes of activation. This finding aligns with theoretical predictions by Boxerman *et al*., who demonstrated that true SEs have the smallest susceptibility difference between deoxygenated and oxygenated blood across all vessel sizes [6]. Consequently, the BOLD CNR induced by the bilateral visual stimulus is notably reduced in SE acquisitions compared to other methods presented in this work, as seen in Fig 2(a).

The reduced BOLD CNR is reflected in the *β*_*contrast*_ terms estimated by the general linear model and the standard deviation of the residuals to the model fit, shown in Fig 2(b) and Fig 2(c). Since t-statistics are proportional to *β*_*contrast*_ and inversely proportional to the standard deviation of the residuals to the model fit, the attenuated BOLD CNR results in fewer voxels exceeding the set significance threshold, leading to sparse activation maps in SE acquisitions shown in Fig 3.

The SE activation findings are consistent with both theoretical and empirical evidence in the literature. Han *et al*. [5] *corroborated Boxerman et al*.’s [6] simulations, emphasizing the need to develop SE acquisitions with improved BOLD sensitivity, even at ultra-high magnetic field strengths such as 9.4 T. The SE activation maps presented in Fig 3 further support these established findings, demonstrating the limited sensitivity of current SE techniques with high-resolution fMRI as performed here.

SE acquisitions can offer an advantage over GRE acquisitions, despite their lower BOLD sensitivity, in some areas of the brain. In the GRE images shown in the first column of Fig 3, substantial signal loss occurs in the cerebral lobules due to susceptibility differences between tissue types, particularly at air-tissue interfaces. This signal loss is also evident in ASE acquisitions in the last two columns of Fig 3. While visual stimulation activation maps, in Fig 3, generated from GRE and ASE data show few active voxels in the cerebral lobules, this could pose challenges for fMRI studies where the evoked response involves brain regions affected by signal loss or in whole brain functional connectivity magnetic resonance imaging (fcMRI) studies. In contrast, SE images demonstrate recovered signal in the cerebral lobules as seen in Fig 3.

SE techniques, while capable of recovering lost signal, are not routinely used in fMRI experiments due to inadequate BOLD sensitivity. However, Boxerman *et al*. have shown that offsetting TE by a duration *τ* can modulate BOLD contrast [6], and others have shown that short spin evolution times can achieve optimal BOLD contrast when such large-scale signal loss is present [21].

The results, presented in Fig 2(a), demonstrate that the BOLD CNR in the ASE acquisitions exceed that of SE acquisitions. This is consistent with the findings in Boxerman *et al*. [6]. The enhanced BOLD sensitivity ASE stems from increasing susceptibility-related *T2*^*′*^ decay from blood susceptibility changes as the time offset *τ* increases. This is reflected in the greater number of active voxels in ASE2 activation maps compared to ASE1 and SE maps (Fig 3).

Theoretically, as *τ* approaches infinity, the BOLD contrast for ASE becomes identical to GRE, suggesting that larger *τ* values should yield greater BOLD contrast. Following this logic, ASE2 should exhibit higher BOLD CNR compared to ASE1. However, this is not evident in Fig 2(a) due to the competing global effect of relaxation reducing the base signal-to-noise ratio. As *τ* increases, signal loss from tissue T2 and T2^*′*^ relaxation begins to dominate, diminishing BOLD sensitivity. This effect is more pronounced at higher magnetic field strengths; at 9.4 T, T2^*′*^ is significantly shorter than at 3 T, causing T2^*′*^ related signal loss to dominate at shorter echo times.

It is worth noting that ASE2 has both increased contrast and noise compared to ASE1 in Fig 2. If the baseline ASE2 images were less noisy with comparable signal to ASE1 images, it is likely that ASE2 would show larger BOLD CNR compared to ASE1. This would be the case if the ASE2 images were measured at a lower field strength or with a shorter TE at 9.4 T. The images in Fig 3 supplements these findings, revealing that ASE2 has more active voxels co-localized with the area of GRE activation.

Furthermore, in S1 Figure, the ASE2 active voxels are shown to be likely true positive results, as they are clustered and the cluster scales in size with significance level. The behavior of ASE techniques observed in Fig 2 and Fig 3 aligns well with the theoretical curves presented by Boxerman *et al*. [6], supporting the validity of our findings.

ASEME-EPI demonstrates both the lowest inter-subject mean *β*_*contrast*_ and standard deviation of the residuals to the fit model in Fig 2(b) and Fig 3(c), respectively. Notably, this does not translate to the lowest inter-subject mean BOLD CNR, Fig 2(a). The key lies in the inherent denoising of the BOLD data using ASEME-EPI, as evidenced by the standard deviation of the residuals which is reduced by more than the contrast.

The denoising effect of ASEME-EPI is attributed to its echo combination method. After T2^*\**^ estimation, echoes are linearly combined into a composite timeseries, with the SE given a weight of 1 and ASEs weighted based on T2^*\**^ relaxation. This approach prioritizes earlier ASEs with higher signal-to-noise ratio, resulting in a composite image primarily composed of high SNR content. Additionally, the combination method exploits the zero-mean characteristic of uncorrelated noise, effectively reducing noise through averaging. This denoising occurs before model fitting, contributing to the low standard deviation of residuals to the model fit using ASEME-EPI, shown in Fig 2(c), and the second-highest number of active voxels in its activation map in Fig 3. While an SNR-optimized combination was used here, other denoising methods will likely yield more striking results.

Kundu *et al*. [22] supplemented the weighted summation method by introducing a denoising technique prior to echo combination. The group noted signal intensity decay seen in multi-echo fMRI (ME-fMRI) studies can be attributed to fluctuations in initial signal intensity *S*_0_ and in transverse relaxation R2^*\**^. Initial signal intensity is dependent upon proton density, T1, and spin excitation history [1],[2],[3]. BOLD-driven changes in blood oxygenation cause a local magnetic field inhomogeneity modulating R2^*\**^, not *S*_0_, and a multi-echo independent component analysis (ME-ICA) method, pioneered by Kundu *et al*. [22], *separates R2*^*\**^ (BOLD) from *S*_0_ (non-BOLD) components. If such a technique were applied to data acquired with the ASEME data, similar reduction in non-BOLD components of noise would be likely. This ME-ICA method, and an echo combination focused on maximizing CNR instead of SNR are both items of future consideration.

As shown in Fig 3, ASEME-EPI activation maps at *α* of 0.01 do not exhibit activations which span anatomical cortical boundaries. GRE requires a much stricter significance level (e.g. *α* = 0.005, S1 Figure) to exclude activations outside V1. This comparison reveals two key points: (1) BOLD contrast from large vessels in GRE produces high t-statistics in voxels, leading to seemingly sensitive and specific responses even at strict thresholds like *α* of 0.01. These are often “large vein effects” rather than true neuronal activations. (2) ASEME-EPI’s lack of blurred activation at *α* of 0.01 results from denoising and preference for higher SNR echoes, a consequence of the combination technique, which attenuates the contribution of larger vessels. Consequently, voxels containing these large vessels are less likely to appear as activations of low anatomical interest.

In addition to the denoising provided by ASEME-EPI, it recovers signal loss compared to a GRE, because it incorporates an SE readout. This results in a combined image that retains signal in the cerebral lobules, as seen in Fig 3, while GRE and ASE cannot recover such signal.

The study described herein is a practical feasibility investigation with a notable limitation of its small sample size. Increasing the sample size in future studies could enable the robust quantitation of differences in CNR, contrast, and noise across the techniques. Additionally, while SE and GRE acquisitions include different sensitivity to the macrovasculature which likely drive differences seen with the ASEME-EPI and GRE acquisitions, a correlative analysis with venography could provide further insight. While the acquisitions included in this work are of exceptionally high spatial resolution, illustrating the benefit of ASEME-EPI compared to SE acquisitions, the low SNR data make an effective quantitative comparison with SE activations fraught. Future work could consider these acquisitions with tasks that include more blocks of stimulus at the expense of longer experimental time (which would be prohibitive with the number of different functional acquisitions considered herein), or lower spatial resolution could be used. Furthermore, increasing the sample size may help inform the wide inter-subject variance observed in this study, as variation of anesthesia with the vasoactive isoflorane agent, used to augment the dexmedetomidine anesthesia.

## Conclusion

This work introduces a new technique to enable the use of multi-echo acquisitions for pre-clinical fMRI at high fields. The ASEME-EPI experiment employs an SE readout followed by two ASE GRE readouts. In the case of high fields, where T2^*\**^ decay has been significantly shortened compared to 3 T, this pulse configuration provides an initial SE which is only T2 attenuated followed by T2^*\**^ attenuated GRE echoes. This configuration creates a higher SNR on later echoes compared to multi-echo GRE or SAGE configurations. In a preliminary exemplar experiment, this ASEME-EPI acquisition yielded BOLD CNR which is not statistically different from GRE, the preeminent choice for fMRI studies conducted both in human and in pre-clinical subjects. Denoising through SNR-optimized averaging echo images, a hallmark of ME-fMRI acquisitions, drove this result. Furthermore, ASEME-EPI recovers signal loss in areas of significant field inhomogeneity, evident in GRE acquisitions, which can be problematic if activation is expected in brain regions with susceptibility-related signal loss. This study lays the foundation for ASEME-EPI and the potential use in BOLD fMRI experiments in pre-clinical subjects at high-field.

## Supporting information

**S1 Appendix. Signal Intensity Derivation**. Signal intensity comparison between ASEME-EPI and GRE-EPI.

**S1 Table. Empirically determined** *α***’s (***α*_*empirical*_**) for various family-wise error (FWE) rates**. The *α*_*empirical*_’s were calculated in a single subject. Each *α*_*empirical*_ is rounded to three decimal places for improved readability.

**S2 Table. Empirically determined** *α***’s (***α*_*empirical*_**) for a set family-wise error (FWE) rate equal to 0.01**. The number of active voxels exceeding *α*_*empirical*_ in the brain is reported in parentheses. Each *α*_*empirical*_ is rounded to three decimal places for improved readability.

**S1 Figure. Single subject, single slice activation maps**. The t-statistic thresholds correspond to the *α*_*empirical*_ values cited in S2 Table. Marked voxels are those that exceeded the t-statistic threshold, and are considered active according to the general linear model fit. The various family-wise error (FWE) rates are shown as rows, and are arranged from top-to-bottom as follows: 0.0005, 0.001, 0.005, 0.01, and 0.05. The different acquisitions are shown as columns, and are arranged from left-to-right as follows: GRE, ASEME, SE16, SE40, SEASE, ASE1, and ASE2.

## Acknowledgments

The animals and visual stimulation used in this publication was supported by the National Eye Institute of the National Institutes of Health under award number U24EY029891.

The authors would like to thank David Schwabe, Matthew Runquist, and Kathleen Yin for their assistance with animal care and handling while in the magnetic environment.

## References

1. Buxton R. The physics of functional magnetic resonance imaging (fMRI). Rep Prog Phys. 2013;76:096601.

2. Haacke E, Brown R, Thompson M, Venkatesan R. Magnetic resonance imaging: physical principles and sequence design. John Wiley & Sons, Inc.; 1999.

3. Kim S, Bandettini P. In: Principles of BOLD Functional MRI. Springer International Publishing; 2023. p. 461–472. Available from: 10.1007/978-3-031-10909-6_19.

4. Kim S. Biophysics of BOLD fMRI investigated with animal models. J Magn Reson. 2018;292:82–89.

5. Han S, Eun S, Cho H, Uludağ K, Kim S. Improvement of sensitivity and specificity for laminar BOLD fMRI with double spin-echo EPI in humans at 7T. Neuroimage. 2021;241:118435.

6. Boxerman J, Hamberg L, Rosen B, Weisskoff R. MR Contrast due to Intravascular Magnetic Susceptibility Perturbations. Magn Reson Med. 1995;34:555–566.

7. Menon R. The great brain versus vein debate. NeuroImage. 2012;62:970–974.

8. Stables L, Kennan R, Jc G. Asymmetric spin-echo imaging of magnetically inhomogeneous systems: Theory, Experiment, and Numerical Systems. Magn Reson in Med. 1998;40:432–442.

9. Posse S, Wiese S, Gembris D, Mathiak K, Kessler C, Grosse-Ruyken M, et al. Enhancement of BOLD-contrast sensitivity by single-shot multi-echo functional MR imaging. Magn Reson Med. 1999;42:87–97.

10. Zhao L, Raithel C, Tisdall M, Detre J, Gottfried J. Leveraging multi-echo EPI to enhance BOLD sensitivity in task-based olfactory fMRI. bioRxiv. 2024;.

11. Gati J, Menon R, Uğurbil K, Rutt B. Experimental determination of the BOLD field strength dependence in vessels and tissues. Magn Reson Med. 1997;38:296–302.

12. Jesmanowicz A, Bandettini P, Hyde J. Single-shot half k-space high-resolution gradient-recalled EPI for fMRI at 3 Tesla. Magn Reson Med. 1998;40:754–762.

13. Schmiedeskamp H, Straka M, Newbould R, Zaharchuk G, Andre J, Olivot J, et al. Combined spin- and gradient-echo perfusion imaging. Magn Reson Med. 2012;68:30–40.

14. Keeling E, Bergamino M, Ragunathan S, Quarles C, Newton A, Stokes A. Optimization and validation of multi-echo, multi-contrast SAGE acquisition in fMRI. Imaging Neuroscience. 2024;.

15. Sidabras J, Mett R, Hyde J. MRI surface coil pair with strong inductive coupling. Rev Sci Instrum. 2016;87.

16. Woolrich M, Beckmann C, Nichols T, Smith S. 7. In: Statistical analysis of fmri data. vol. 19. Neuromethods: New York, New York: Spring Science and Business Media; 2016. p. 183–239.

17. Birn R, Smith M, Jones T, Bandettini P. The respiration response function: the temporal dynamics of fMRI signal fluctuations related to changes in respiration. Neuroimage. 2008;40:644–654.

18. Menon R. Postacquisition suppression of large-vessel BOLD signals in high-resolution fMRI. Magn Reson Med. 2002;47:1–9.

19. Nencka A, Rowe D. Reducing the unwanted draining vein BOLD contribution in fMRI with statistical post-processing methods. Neuroimage. 2007;37:178–188.

20. Zhou J, Ni R. The Tree Shrew (Tupaia belangeri chinensis) Brain in Stereotaxic Coordinates. Springer Singapore; 2016.

21. Miyapuram K, Noortlan O, Tobler P, Schwarzbauer C. Imaging Brain Regions with Susceptibility-induced Signal Losses using Gradient and Spin Echo Techniques. Proceedings of the 31st Annual Meeting of the Cognitive Science Society. 2007;.

22. Kundu P, Inati S, Evans J, Luh W, Bandettini P. Differentiating BOLD and non-BOLD signals in fMRI time series using multi-echo EPI. NeuroImage. 2012;60:1759–1770.

